# The influence of environment on mosquito feeding patterns: a meta-analysis of ‘universal’ DNA diet studies in a global context

**DOI:** 10.1101/2024.02.21.581447

**Authors:** Richard O′Rorke, Meshach Lee, Nicholas J. Clark, Tamsyn Uren Webster, Konstans Wells

## Abstract

Mosquitoes have innate preferences for their blood-meal hosts, but these can be modified by the environment, with implications for disease spread under climate and land use change. To predict the spread of vector-borne pathogens more accurately we need to better understand blood-meal plasticity under changing environmental conditions. We compiled blood-meal studies for six prominent disease vectoring mosquitoes from around the globe (*Aedes aegypti, Ae. albopictus*, *Anopheles funestus, An. gambiae*, *Cx. pipiens, and Cx. quinquefasciatus*). We targeted studies that used universal molecular methods (typically PCR/metabarcoding) to identify hosts from a wide range of candidates (as opposed to studies using methods that presuppose host identity - such as precipitin, ELISA). We found that blood-meals from the >15,600 analysed were mostly from the expected host-groups for each mosquito species, but we frequently encountered atypical hosts (e.g. mammalophilic species feeding on birds/reptiles). The universal methods used by the studies in our metanaalysis identified high host richness, and we found ≥174 hosts for *Culex* and ≤65 species for *Aedes* mosquitoes at a considerably increased discovery rate of novel hosts per sampling effort. We used a hierarchical Dirichlet regression model to analyse global variation in feeding patterns in relation to environmental datasets (land use, precipitation, mean annual temperature, latitude, human and livestock density). Land use, mean annual temperature and poultry density had noticeable effects on blood-meal selection of *Ae. aegypti*, *Cx. pipiens* and *Cx. quinquefasciatus*. Although host density was a factor in blood-meal selection - host choice is not a simple function of host availability, as has previously been observed, but contingent on other drivers. While our compiled dataset afforded us these insights, improving resolution and consistency of data gathering and reporting would improve the precision of how blood-meal studies can inform us of present and potential risks of pathogen transmission events.

## 1 INTRODUCTION

The contingency of female mosquito blood-feeding patterns on the environment is likely to have consequences for the transmission of vector-borne pathogens. For example, increased preference for human blood-meals can increase the transmission of pathogens between humans, while mixed host utilisation increases the risk for zoonotic pathogen transmission (Fikrig & Harrington, 2021). With a global rise in arboviral spread (Messina et al., 2015; Weaver et al., 2018) understanding how changing environmental conditions affect blood-feeding patterns would improve prediction of pathogen spillover.

Generally, certain mosquito species have innate preferences for certain hosts. Some mosquitos, such as *Anopheles gambiae* and *Aedes aegypti*, are considered to exhibit strong and consistent preferences for feeding on humans in both laboratory and field contexts (McBride, 2016; Richards et al., 2006).

However, other mosquito blood feeding patterns are synergistically modified by additional intrinsic and extrinsic factors such as host seeking cues and host detection, behavioural plasticity, host availability, and pathogen infection (Harrington et al., 2014; Lyimo & Ferguson, 2009; Yan et al., 2021). For example, *An. arabiensis* feeds relatively more on livestock than humans with increasing livestock density (Mayagaya et al., 2015). Host availability is generally considered a strong environmental factor impacting host selection of some mosquito species, with forage ratio (i.e. the ratio of actual blood-meals over available host species) studies being a prevailing approach since the late 1960’s. However, these forage ratio studies consistently reveal there are more factors determining host selection than just host availability (Riccetti et al., 2022; Yan et al., 2021). Such variability in host selection appears to be species- or even biotype-specific; for example, some forms of the *Culex pipiens* species complex are considered bird specialists, but readily generalise and some forms appear to predominantly feed on mammals (Fonseca et al., 2004).

Changing climate and land use modify the distribution of vectors and hosts (Altizer et al., 2013). Under very dry conditions, for example, artificial water bodies in more urban environment may offer the most resilient breeding environments (Rose et al., 2020). Also, global-scale changes modify vector phenotypes and the complex interactions that determine host choice. Nutrient or temperature-induced stress can change the frequency and selectivity of mosquito host seeking behaviour, as has been evidenced in laboratory studies (reviewed in (Carvajal-Lago et al., 2021)). To better understand these dynamics, we can take advantage of the freely available and independently collected global environmental datasets to identify global-scale patterns, particularly in the context of a changing environment.

A hundred years ago, (King & Bull, 1923) published their study that used a molecular antigen-binding technique (precipitin) to determine the origins of blood-meals recovered from guts of engorged female mosquitos captured from the wild. Numerous studies have since used various molecular techniques to identify what species mosquitoes are feeding on. Some of these are very targeted (identifying one species such as human or cattle) and other molecular methods can identify a broad range of hosts (reviewed in (Apperson et al., 2002; Mukabana et al., 2002; Takken & Verhulst, 2013). The advantages of using such broad approaches, typically referred to as “universal”, is that they do not presuppose the identity of the blood-meal host of the mosquitoes, and allow more accurate identification of host range and foraging ratios (Ferraguti et al., 2021; Hernández-Andrade et al., 2019; Melgarejo-Colmenares et al., 2022). Universal approaches have also recovered exciting novel information on the unexpected diversity of mosquito’s host ranges including the importance of amphibians, reptiles, fish and even some invertebrates (Börstler et al., 2016; Harrington et al., 2014; Molaei et al., 2006). Universal studies constitute a very small percentage of the host diet studies conducted thus far (especially for taxa such as *An. funestus, An. gambiae* and *An. arabiensis*) but are expanding our insights into the plasticity of host preferences (e.g. (Reeves & Burkett-Cadena, 2022)) and have the potential to inform us of the local transmission risk of different pathogens (Hernández-Andrade et al., 2019).

Most meta-analyses on mosquito feeding patterns conducted so far have combined universal and species-specific methods that target expected/known hosts (especially humans and livestock) and risk missing out on wider host shifts that might be expected under global change scenarios (Cebrián-Camisón et al., 2020; Chaves et al., 2010; Stephenson et al., 2019). Here, we compile studies based on universal PCR methods to explore how gradients in host densities, climate, and land use influences mosquito blood-feeding patterns. This quantitative meta-analysis focuses on compositional data compiled on the realized host utilisation of six common mosquito species that pose zoonotic disease risks and exhibit various degrees host feeding fidelity. Our approach, although correlative (rather than mechanistic), offers new perspectives on vector feeding behaviour and therefore what drives vector-mediated disease transmission in a globally changing environment.

## 2 MATERIALS AND METHODS

### 2.1 Focal species and blood-meal data extraction

We focused on six mosquito species: *Aedes aegypti*, *Aedes albopictus*, *Anopheles gambiae*, *Anopheles funestus*, *Culex pipiens* s.l., and *Culex quinquefasciatus*. These are of considerable interest as vectors of infectious diseases and feed to various extents on humans (Chaves et al., 2010; Takken & Verhulst, 2013). We conducted two systematic literature searches that reported counts of mosquitos feeding on different blood-meal hosts. Our initial search (August 2022) was broad and afforded us a “bird’s eye view” of the literature. For this, we used Web of Science, Pubmed, Scopus, GoogleScholar online databases with combinations of the search terms ‘bloodmeal*’, ‘blood meal’, blood-meal’, ‘feeding’, ‘habit’, ‘pattern*’, ‘preference*’, ‘interaction*’, combined with ‘mosquito*’, ‘vector*’, ‘vector-host’, ‘host*’, ‘vertebrate*’, ‘animal*’. The asterisk (‘*’) was used to search for possible variations of terms. Our subsequent systematic search of the literature (July-September 2023) focussed on Web-of-Science and involved two approaches. The first used the phrase ‘bloodmeal*’ OR ‘blood meal’ OR ’blood-meal’” and the names (including synonyms) of the focal mosquito species. If the search returned too many irrelevant hits (>200), then additional ‘methods’ search terms were used (**Table S1**). Another search used publications of popular PCR primer sets as a search term (**Table S1**). If the search returned too many irrelevant hits, then the number of linked articles was refined using the search term ‘mosquito*’ (**Table S1**). This resulted in 1,495 published articles that we subsequently filtered with the following criteria:

1) The arthropod vector was one of six focal species
2) PCR primers used to identify blood-meal hosts were ‘universal’ (for vertebrates/mammals/birds) and not targeted to particular species (Table S1b)
3) Either Sanger (dideoxy) sequencing or hight-throughput DNA sequencing is used at some point to confirm host identity
4) Spatial information was provided as either geographical coordinates or salient location names and sampling sites were not pooled over spatial extents >50km in diameter
5) Host species identity data was available
6) For any given mosquito species and sampling site, there must be ≥7 bloodmeals (corresponding to an ∼80% chance of detecting an uncommon host present in ∼20% of samples).

This resulted in 88 studies that covered 121 different study sites (some studies provided data for multiple sites, see **Figure S1** for location map). From these we generated a database of geographical coordinates, counts of blood-meals and the host species (resolved to smallest taxonomic unit) for each mosquito species. Counts of blood-meal origins were assigned to both hosts in the sixteen studies that recorded mosquitos feeding on multiple hosts (e.g. ‘human + cow’). All extracted scientific species names were checked and aligned to the Integrated Taxonomic Information System (ITIS) database, using the *taxize* 0.9.1 R package (Chamberlain & Szöcs, 2013).

For studies with missing geographical coordinates, we identified coordinates for location names at smallest available administrative units using various online maps and search engines. For studies with multiple sites/coordinates from a single region (<50 km in diameter), we computed the average of coordinates for use in analyses.

### 2.2 Host functional and taxonomic groups

Host functional groups were derived and consisted of: *f1*) ‘Human’, *f2*) ‘Mammal - pets’ (cat, dog, rabbits if specified as domestic), *f3*) ‘Mammal - farmed’ (cattle, buffalo, donkey, goat, horse, sheep, pigs unless explicitly recorded as wild), *f4*) ‘Mammal - wild’ (all other mammalian species), *f5*) ‘Bird - farmed’ (commensal chickens, goose, ducks if recorded as farmed animals), and *f6*) ‘Bird - wild’ (all free-roaming wild birds). All other origins were grouped as ‘other’ (including zoo animals, caged ornamental birds, reptiles, and amphibians). We also grouped host species in taxonomic groups using different taxonomic levels according to 4% overall sample origins thresholds (these were *‘Homo’, t2) ‘Gallus’, t3) ‘Canidae’, t5) ‘Felidae’, t6) ‘Bovidae’, t7) ‘Mammalia’, t8) ‘Cardinalidae’, t9) ‘Columbidae’, t10) ‘Corvidae‘, t11) ‘Turdidae‘, t12) ‘Aves‘, t13) ‘others’* (amphibian and reptiles).

The total number of recorded blood-meals per vector and the number of sites (*s*) were: *Ae. aegypti*: 1,803 (s=18), *Ae. albopictus*: 1,769 (s=22); *An. funestus*: 308 (s=5); *An. gambiae*: 631 (s=15); Cx. pipiens complex: 11,172 (s=87) with *Cx. pipiens* s.l.: 4,886 (s=48); *Cx. quinquefasciatus*: 5,449 (s=36). Out of the *Cx. pipiens* s.l. samples, 1,061 (s=8) were more specifically assigned to the *Cx. pipiens* f. *pipiens biotype and 45* (s=4) *to the Cx. pipiens* f. *molestus* biotype.

### 2.3 Environmental variables

We selected nine environmental predictor variables (extracted with a 10km radius around each sites’ geographical coordinates) from a larger set of environmental variables (**Table S2**) by removing correlated variables (r ≥ 0.7) and selecting those with strongest correlations to the response variables and by ordination of land cover variables (**Figure S2, S3**). Specifically, the selected environmental predictor variables were: *x.1*) ‘annual mean temperature’ (bio1), *x.2*) ‘annual rainfall’ (bio12), and *x.3*) ‘rainfall of driest month’ (bio14). Climatic variables were obtained from the WorldClim database of gridded climate data at a 0.01-degree resolution ((Fick & Hijmans, 2017), 2017; http://worldclim.org/version2). Land use types were based on Copernicus landcover data from 2010 (map version 2.07; https://cds.climate.copernicus.eu; land cover categories were ‘urban’, ‘water bodies’, ‘cropland’, ‘grassland’, ‘shrubland’, and ‘tree cover’) and the normalized difference vegetation index (NDVI, computed as mean and one SD of all measures values within 10-km buffer) for the year 2010 from the Terra Moderate Resolution Imaging Spectroradiometer (MODIS, MOD13Q1 version 6, https://lpdaac.usgs.gov/products/mod13q1v006/). As land cover variables are typically highly correlated, we used principal component analysis (PCA) to generate two principal components. The first was *x.4*) ‘land cover PC1’, (green space) where increasing values represent mainly increasing vegetated land cover and NDVI and decreasing water bodies and urban land cover. PC1 explained 31.1% of the variation in land cover variables across sites. The second was *x.5*) ‘land cover PC2’ (urban space) where increasing values mainly represent increasing urban land cover and decreasing tree cover and explained 18.4% of land cover variation across sites (**Table S3**). Next, *x.6*) ‘human population density’ was based on the Gridded Population of the World version 4.0 dataset (GPW4) (Center for International Earth Science Information Network, 2018). The *x.7*) ‘ruminant livestock’ density pooled local numbers of sheep, goats, cattle, and buffalo and *x.8*) ‘poultry density’ pooled local numbers of chickens and ducks using the Gridded Livestock of the World (GLW3) in 2010 at approximately 10 km^2^ resolution (Gilbert et al., 2018). Finally, we included *x.9*) latitude as an environmental predictor variable to account for the large-scale variation in biological diversity and other factors along this prominent gradient. While we also considered studies that were conducted indoors (or indoors & outdoors), we did not account for this possible sampling bias as random effects in our analysis because of the small number of indoor studies (≤ 3 per mosquito species); we rather repeated all analysis while excluding indoor studies and present these results as supplementary material.

### 2.4 Statistical analysis

Our key aims were to explore variation in mosquito host utilisation across sites and to identify shifts in host selection associated with environmental predictor variables for each of six focal mosquito species. We implemented the model described below through a Bayesian workflow with different model versions fitted to simulated and empirical data for optimisation. The final model was fitted separately for each mosquito species (and biotype), but for *Culex* mosquitoes, we additionally ran the model for the ‘*Cx. pipiens* complex’ as a whole (to account for studies that did not differentiate between members of the complex).

Dirichlet regression was used because it allows us to model the relationship between the parameters representing the proportions of distinct host categories/taxa and environmental predictor variables, resembling data structures commonly analysed in diet and microbiome compositional studies (Douma & Weedon, 2019). We combined the Dirichlet model in a hierarchical multivariate modelling framework with a Binomial model of the counts of blood-meals originating from different host groups to account for uncertainty in proportions arising from small and unequal sample sizes. We modelled the proportions of blood-meals θ_*g,s*_ that originated from any host group *g* out of G=7 at site *s* based on the number of mosquitos *n_g,s_* with blood-meals reported from that group and the total number of mosquitos *N_s_* for which blood-meals were identified at that site as:

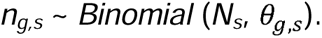

Here, θ_*g,s*_ is assumed to be a *G*-simplex *Θ_s_* of the interlaced feeding proportions of the four different host groups *Θ_s_* subject to the constraint that proportions sum to one.

We therefore modelled

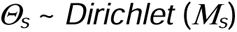

whereby the vector *M_s_* comprises a normalized exponential function of the four joint linear predictors terms μ_*g,s*_:

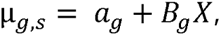

where *a*_*g*_ is the group-specific intercept, *X* is a matrix of the environmental predictor variables described above and *Β* is a vector of coefficient estimates for the predictors denoted in the matrix *X*.

We fitted models for parameter estimation in Stan (Carpenter et al., 2017; Team, 2023) via the *CmdStanr* interface (https://mc-stan.org/cmdstanr/). Stan’s probabilistic modelling framework based on Hamiltonian Monte Carlo enables efficient Bayesian workflows for model specification and diagnostics and exploring the uncertainty in estimates for challenging problems (Betancourt, 2017). Using this sampling approach, we randomly imputed missing covariate data (in GLW3 and GPW4) by averaging over the uncertainty in their values (randomly drawn from *Normal*(μ=0, σ=1) distributions for variables scaled to have variance equal to 1).

Since feature selection and parameter estimation in multiple regression models can be challenged by high-dimensional feature space for sparse data (Piironen et al., 2020) we tested *Normal*(μ=0, σ=1), *Student.t*(ν=3, μ=0, σ=1), and regularised horseshoe (RHS) priors with various global and local shrinkage priors (Piironen & Vehtari, 2017) for the coefficient estimates of *Β*. We then iteratively fitted the model to simulated and real data with each of four chains sampling 1,000 posterior draws after warmups of 10,000 iterations for optimizing prior specifications. We used trace plots and rank normalized split-R^ for convergence diagnostics and computed the sum of squared residuals for comparing model fits with different priors. We scaled all covariates (centred around zero and variance equal to one) for parameter estimation. Model fit was similar for all three prior specifications, while the RHS priors shrunk more posterior distributions towards zero; we therefore reported results from models with RHS only as the most conservative approach. We assumed those coefficients for which estimates did not overlap zero based on 90% highest posterior density credible intervals (CI) represent meaningful trends and report these as the odds ratios (OR) from the scaled covariates. We used posterior estimates of the model intercepts *a_g_* to draw Dirichlet posterior estimates of the ‘global’ average feeding proportions Ψ_*g*_ and interpreted upper/lower CIs as maximum/minimum feeding proportions on different host groups. We calculated species richness estimates and rarefaction curves based on Hill numbers in the packages iNEXT (Hsieh et al., 2016), using the cumulative number of blood-meal counts from different host species for *Aedes* and *Culex* species. For species richness estimates, we also considered studies with <7 total blood-meals reported (totalling n= 101 studies) in order to maximize the evidence of known host ranges. We used R v4.3.0 (R Development Core Team, 2023) for all analyses and graphics.

## 3 RESULTS

### 3.1 Global scale feeding patterns

Across all sites recovered in the metanalysis the feeding patterns of all focal mosquito species varied in their utilisation of human, other mammalian and avian species (and occasionally amphibians or reptiles - **Figure 1**). The ‘global’ average proportions of blood-meals from humans, according to global Ψ*_human_* estimates, were highest for *Aedes aegyti* (Ψ*_human_*CI of 41 – 95%) and *Anopheles gambiae* (Ψ*_human_*CI of 24-98%). The large credible intervals in these estimates were partly explained by the strong variation in feeding patterns across sites. For *Ae. aegypti*, for example, >80% of blood-meals originated from humans at 4 out of 18 sites (according to θ*_human_*lower CIs) but <40% of blood-meals originated from humans at another 3 sites (according to θ*_human_* upper Cis, **Figure 2**). The *Culex* species in this study were clearly identified as ornithophilic and fed frequently on birds but less often on humans (all Ψ*_human_* CIs 1 – 86%) than the *Aedes* and *Anopheles* species (**Figure 1**).

**Figure 1.**
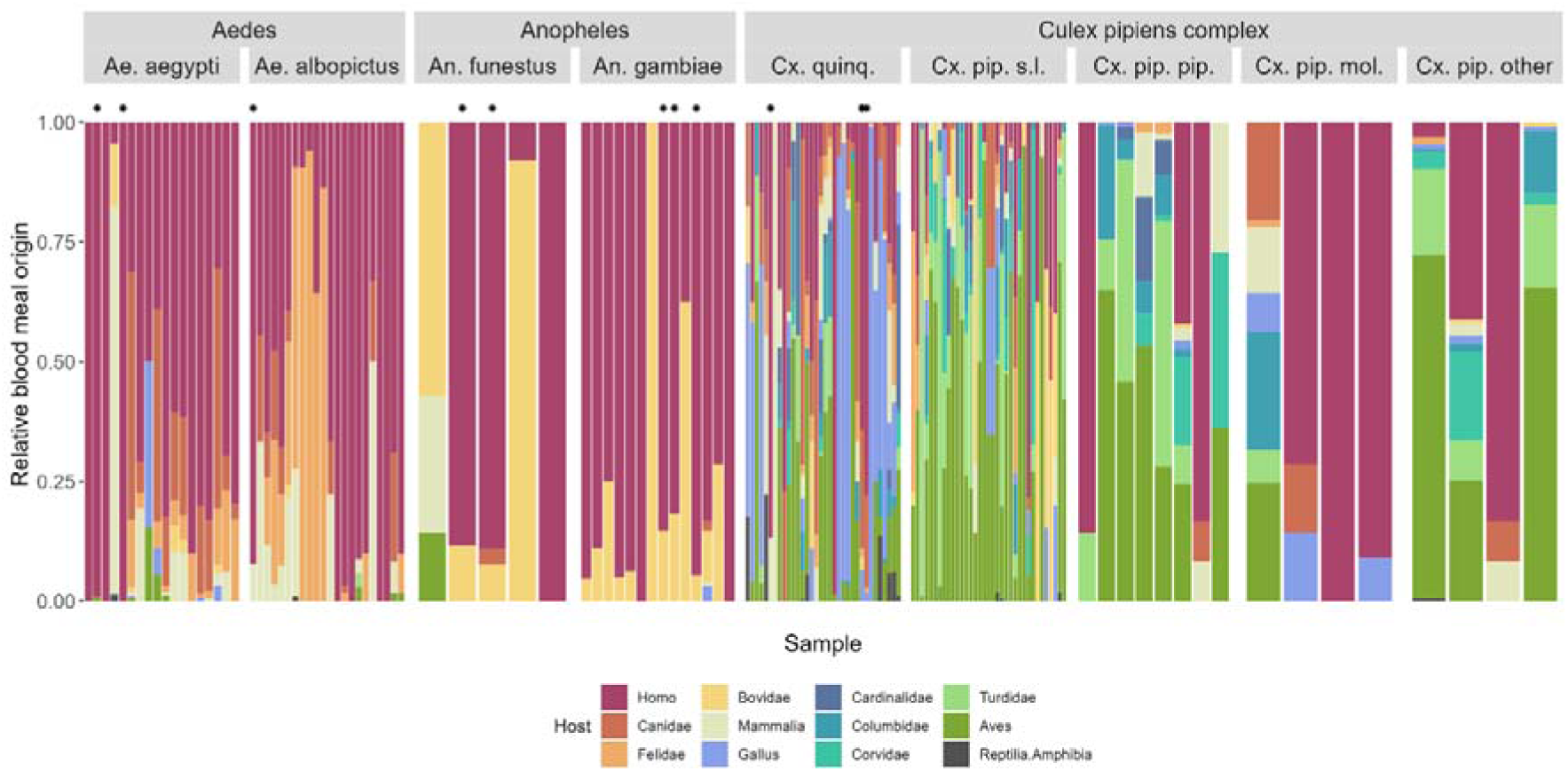
Taxonomic composition of mosquito blood-meal origins from vertebrate hosts plotted as the relative proportion of blood-meals recorded from each host groups at different sampling sites for the focal mosquito species. The plot is faceted by mosquito species with *Aedes* and *Anopheles* species on the left and members of the *Culex pipiens* complex on the right. The latter is comprised of *Cx. quinquefasciatus* and *Cx. pipiens* sensu lato, which is dived into the biotypes *Cx. pipiens f. molestus* and *Cx. pipiens f. pipiens* where known. The ‘Cx. pip. other’ category refers to other specimens that are part of the complex or, more often, studies where members of the complex could not be determined. Asterisks mark those samples from studies conducted indoors only, as one of a number of possible sampling biases in data compiled from multiple studies.

**Figure 2.**
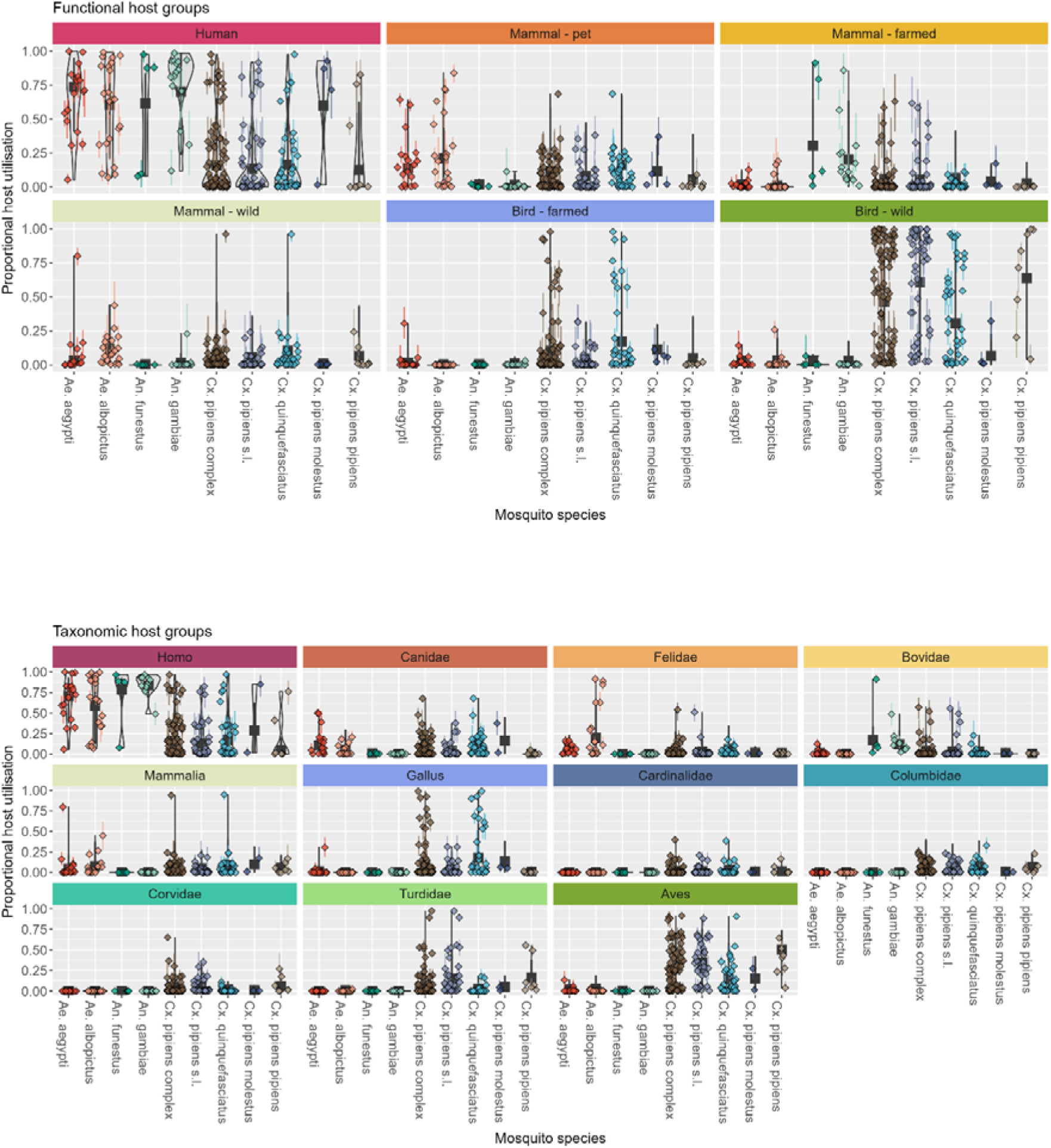
Variation in host feeding patterns of six common mosquito species reported from blood-meal studies at different sites. Coloured point clusters within violins show the relative host utilisation frequencies of A) different functional host groups (humans, domestic and wildlife species) and B) different taxonomic host groups (genera/families/classes of mammals and birds according to relative abundance in the overall data set) at different study sites, with vertical error bars around points representing 95% highest posterior credible intervals of the probabilistic frequencies linked to data observations. Black squares and error bars represent the ‘average’ posterior estimates based on a hierarchical hyperprior modelling approach. Abbreviated genera names correspond to *Aedes, Anopheles and Culex*. *Cx. pipiens* s.l. and *Cx. quinquefasciatus* samples were also combined into *Cx. pipiens complex* and we also conducted analysis for *Cx. pipiens* f. *pipiens and Cx. pipiens* f. *molestus* biotypes.

Functional group utilisation clearly differed among mosquito genera (**Figure 2a**). In most cases, *Aedes* fed predominantly on humans but also fed frequently on mammalian pets, including Canidae and Felidae (≥30% of blood-meals originated from mammalian pets at 3 out of 18 and 6 out of 22 study sites for *Ae. aegypti* and *Ae. albopictus, respectively*) and to a lesser extent wild mammals. *Anopheles* species, in turn, also deviated occasionally from humans and shifted towards feeding on farmed mammals, including Bovidae (**Figure 2**). *Aedes* fed occasionally on wild birds or poultry.

Despite their ornithophilic categorisation, *Culex* species regularly fed on wild, farmed and pet mammals (**Figure 2**). *Cx. quinquefasciatus* fed frequently on farmed birds including domestic chicken *Gallus* (≥60% meals from farmed birds at 6 out of 36 sites), whereas *Cx. pipiens* s.l. predominantly fed on farmed birds (≤44% at all 48 study sites **Figure 2**). *Cx. pipiens* f. *pipiens* fed ≥60% on wild birds at 4 out of 8 sites and ≥60% on humans at another site, whereas *Cx. pipiens* f. *molestus* fed at 2 out of 4 sites ≥60% on humans (according θ estimates), providing insufficient statistical evidence to infer whether these two biotypes differ in terms of mammalophilic feeding preferences.

### 3.2 Shifts in feeding patterns across environmental gradients

We found some signals of environmental conditions predicting changes in feeding patterns across sites (**Figure 3**), despite the overall sparse data for some species due to the inclusion criteria. For *Ae. aegypti*, more blood-meals originated from humans with increasing poultry density (OR: 1.32 – 4.21; but see Figure S3: this effect vanished when excluding indoor studies). More blood-meals of *Ae. aegypti* originated from wild mammals with higher ruminant density (OR: 1.52 – 5.22) and more originated from mammalian pets with higher mean temperatures (OR: 1.06 – 3.68). The model gave concordant results when run with either taxonomic or functionally clustered data.

**Figure 3.**
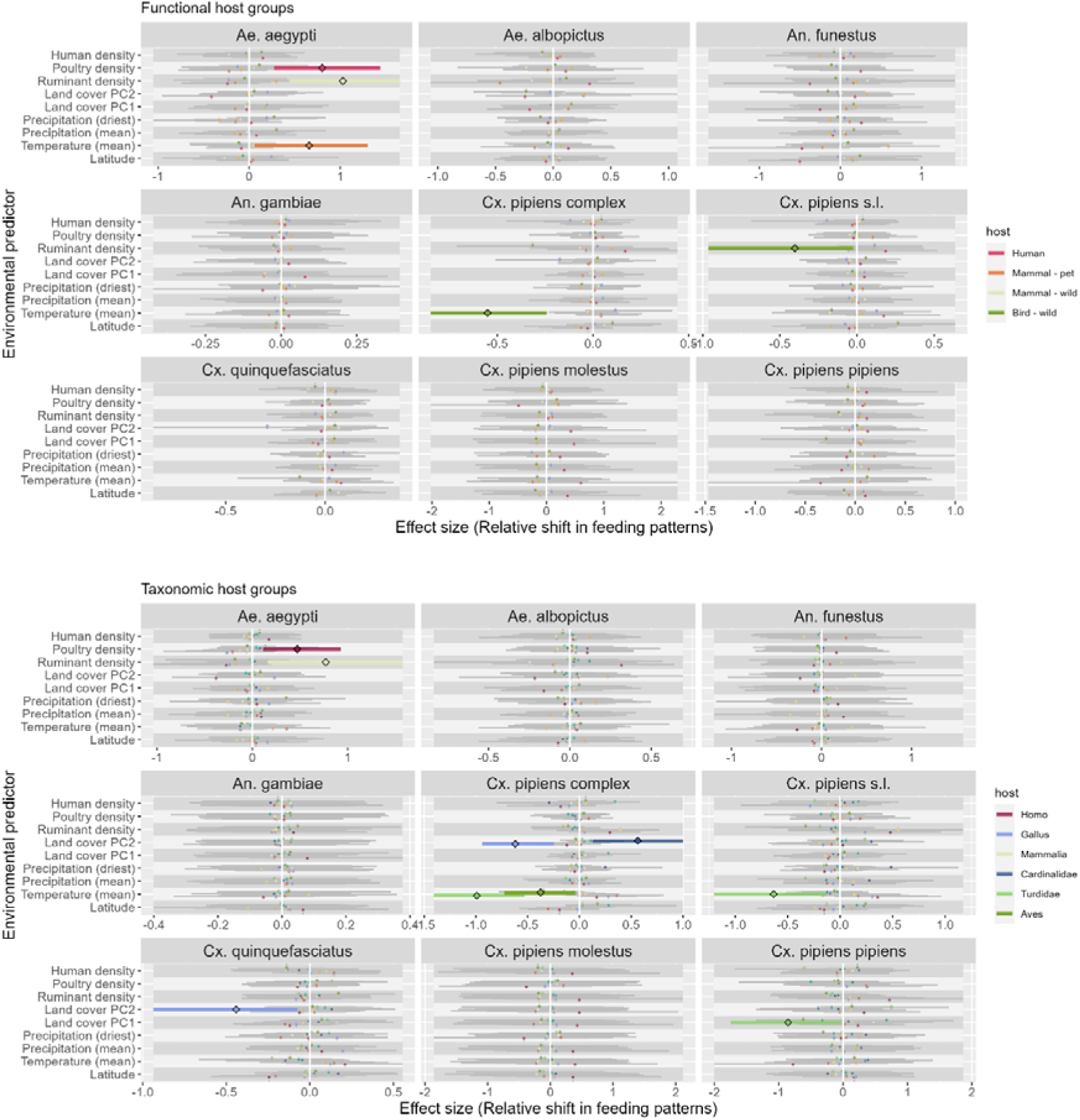
Relative effects of different environmental predictors on the shifts in mosquito blood-meal origin from different functional (panel A) and taxonomic (panel B) host groups. Points and error bars represent the scaled coefficient estimates (mean and 90% CI credible intervals) from models of variation in relative host utilisation frequencies for humans (red), mammalian pets (blue), mammalian wildlife (green) and wild birds (dark blue), whereby only estimates not overlapping one are displayed in thick-lined colour and those overlapping one in thin-lined grey (including all host groups for which no relationship were evidenced). Abbreviated genera names correspond to *Aedes, Anopheles and Culex*. *Cx. pipiens* s.l. and *Cx. quinquefasciatus* samples were also combined into *Cx. pipiens complex* and we also conducted analysis for *Cx. pipiens* f*. pipiens and Cx. pipiens* f*. molestus* biotypes. Higher “Land cover PC1” correlates with higher green vegetated spaces and “Land PC2” correlates with increased urban and decreased tree cover.

Mosquitoes from the *Cx. pipiens* complex fed less often on wild birds when under higher temperatures (OR: 0.43 – 0.79), whereby taxonomic grouping more specifically identified that *Cx. pipiens* s.l. fed less often on Turdidae with increasing temperature (OR: 0.43 – 0.79). With increasing ruminant density *Cx. pipiens* s.l. fed less often on wild birds (OR: 0.38 – 0.98, **Figure 3**). With more urban space (land cover PC2), less blood-meals of *Cx. quinquefasciatus* originated from domestic chicken (OR: 0.34 – 0.90). In urban spaces *Cx. pipiens* complex fed less on domestic chickens and more on Cardinalidae (**Figure 3**). With more green space (land cover PC1), blood-meals of *Cx. pipiens f. pipiens* were less often from Turdidae (OR: 0.17 – 0.97, **Figure 3**). Most of these effects were similar when removing indoor studies from the analysis, while some effects were less evident or even disappeared (**Figure S4**), highlighting the uncertainty arising from possible sampling bias.

### 3.3 Host ranges of Culex are considerably larger than for Aedes species

*Culex quinquefasciatus* had the largest host range of 170 reported host species from 70 families (including 130 bird and 33 mammalian species), followed by *Cx. pipiens* s.l which had 138 host species from 62 different families (including 115 bird and 20 mammalian species). A total of 25 host species from 21 families (including 9 bird and 15 mammalian species) were reported for *Ae. aegypti and 29* host species from 25 families (including 7 bird and 19 mammalian species) for *Ae. albopictus*. *Anopheles funestus* reported 7 host species (including the wild bird *Crinifer piscator*) and *An. gambiae* had 8 host species or (including domestic chicken and the rodent *Thryonomys swinderianus*). Host species richness estimates were 242 (95% CI: 207 - 309) for *Cx. quinquefasciatus*, 225 (95% CI: 179 - 321) *Cx. pipiens* s.l., and only 28 (95% CI: 24 - 45) for *Ae. aegypti* and 37 (95% CI: 30 - 65) for *Ae. albopictus*, respectively. Rarefaction curves indicate that for each genera of *Aedes* and *Culex* there is a particular and distinct host-species discovery rate (**Figure 4**).

**Figure 4.**
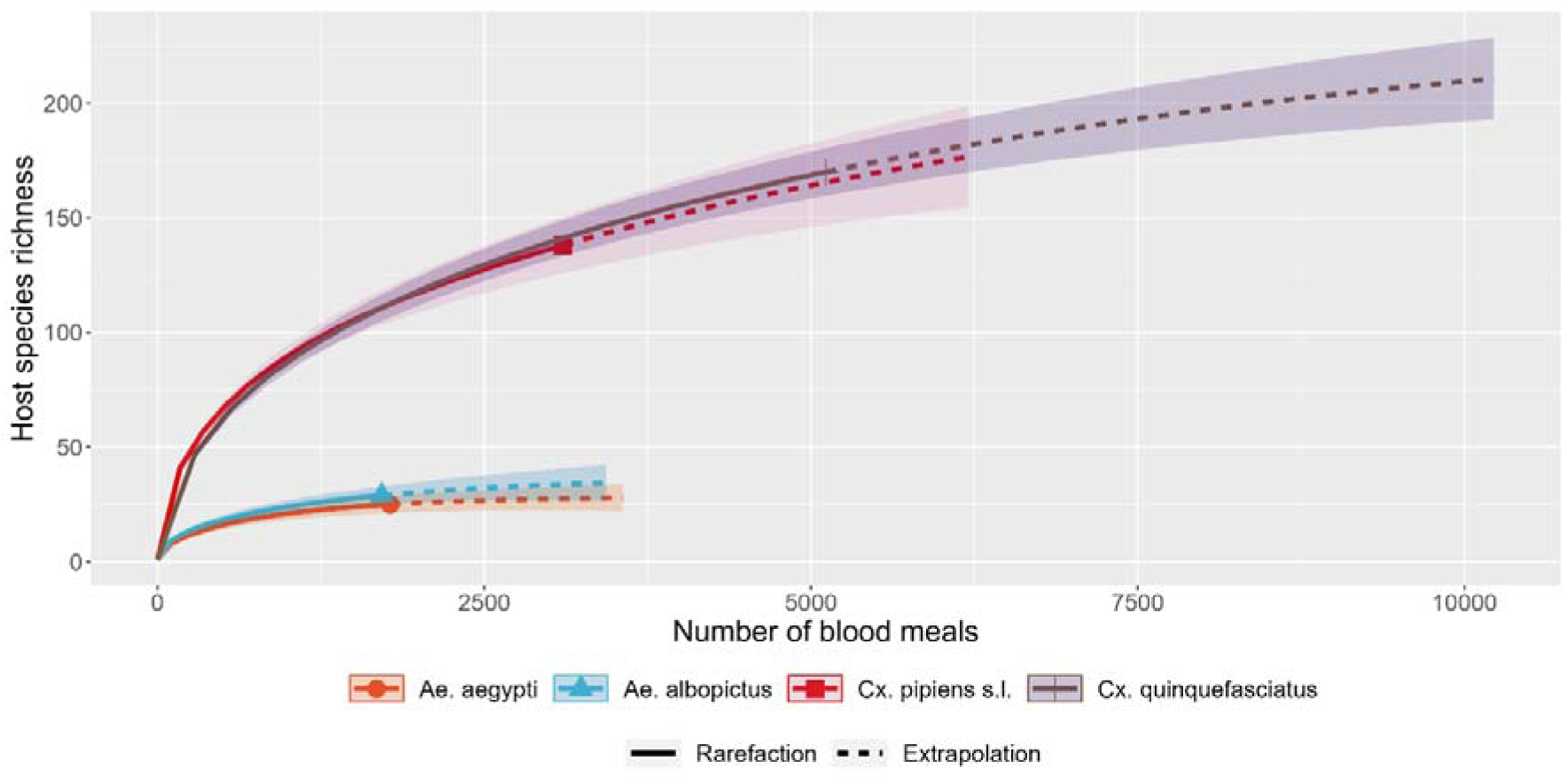
Rarefied host species richness estimates and predictions based on the cumulative number of host species recorded in blood-meal studies for common mosquito species. Estimates (lines) and 95% confidence intervals (shaded areas) are based on Hill numbers. Solid lines refer to the rarefied samples, dashed lines to extrapolated predictions of the same number of samples. Abbreviated genera names correspond to *Aedes and Culex*.

## 4 DISCUSSION

### 4.1 Host range

Our meta-analysis revealed a total of 341 host species across all focal vector species (avian: 267, mammal: 60, reptile: 13, amphibian: 1). The discovery of novel hosts per sampling effort (**Figure 4**) was also at more than double the rate found in other meta-analyses (Bellekom et al., 2021) because we focussed on just papers that used “universal” detection methods (i.e., accumulation curves made with methods that limit the number of hosts discoverable will always tend to plateau before methods that don’t have such limits). This focus builds our understanding of ectoparasite host selection because universal methods are less likely to underestimate the actual range of hosts utilised in the wild. The greatest host richness was witnessed in *Cx. pipiens* s.l. and *Cx. quinquefasciatus*, which had host range estimates of 179 - 321 and 207 - 309 host species respectively. *Ae. aegypti* and *Ae. albopictus* also had high diversity (host range estimates of 24 - 45 and 30 - 65), though considerably lower than *Culex.* This accords with previous studies (Martinet et al., 2019) and most probably reflects the higher diversity of bird species across global habitats (Barrowclough et al., 2016) and that these *Culex* species tend to be ornithophilic whereas *Aedes* are mammalophilic.

#### 4.1.2 The innate vs. realised host niche

Many studies have established that mosquito species have innate host preferences (Lyimo & Ferguson, 2009; Main et al., 2016) that vary from strong to weak (e.g. (Mayagaya et al., 2015)). Although variations can be resolved with lab-based behavioural studies the permutations of host species and environmental contingencies is immense (Fikrig & Harrington, 2021), which is why we also need to look at field studies. We found that the host blood-meals of vectors was according to “type”: e.g. *Anopheles* fed predominantly on mammals. However, there are a few reports of *Anopheles* feeding on birds (Diallo et al., 2013; Ogola et al., 2017; Omondi et al., 2015). Notably, these studies looked at more diverse landscapes and host species assemblages than had previously been considered for this genus (**Figure 1**), showing that although only a few *Anopheles* blood-meals have been studied with universal primers that there is more to discover about their diet selection. For *Aedes*, host-composition varies considerably within mammals (figure 1 and 2), which is broadly consistent with previous observations and possibly has an environmental explanation (e.g. (Little et al., 2022)). This reinforces the caution recommended in assuming *Aedes* species are highly anthropophilic (Bellekom et al., 2021; Ponlawat & Harrington, 2005). Notably, we found that a few taxa dominate, with humans and cats (e.g. (Little et al., 2021; Ogola et al., 2017)) constituting large proportions of hosts of *Ae. albopictus* whereas *Ae. aegypti* studies predominantly reported meals on humans and, to a lesser extent, dogs (e.g. (Estrada-Franco et al., 2021; Hopken et al., 2021)). We found no strong statistical evidence for the recently debated (Börstler et al., 2016; Farajollahi et al., 2011; Fonseca et al., 2004) assumption that *Cx. pipiens* f. *molestus* is mammalophilic and exhibits a distinct host preference from the ornithophilic feeding pattern of *Cx. pipiens* f. *pipiens*.

### 4.2 Global correlation of environmental and host change

We used standardized global datasets for environmental conditions as predictor variables to understand host-feeding patterns of six mosquito species in a hierarchical Dirichlet regression model.

#### 4.2.1 Host density

There were several cases where the feeding selection of mosquitoes on a given host was correlated with the density of another vertebrate (**Figure 3**). This pattern has been observed in previous forage ratio studies and suggests host choice is not a simple function of host availability, but contingent on other unmeasured drivers (Riccetti et al., 2022; Yan et al., 2021). However, we can make fairly good conjectures on what those drivers are. For example, livestock were associated with increased feeding rates of *Ae. aegypti* on wild mammals, but not farmed animals *per se*. This could be due to farms creating conditions that are favourable for a select subset of commensal mammals to proliferate, such as rodents and rabbits. The inverse correlation of *Cx. pipiens* s.l. feeding on wild birds and ruminant density is most likely due to removal of bird habitat on farms. Where poultry density increased *Ae. aegypti* feeding rates were higher on humans, but the ‘human density’ covariate had negligible explanatory effect – this is more difficult to explain but could be a sampling artefact (see supplementary figure S3) or, if not, it could be because the presence of chickens is a proxy for suburbia and farms where moderate human densities coincide with suitable mosquito habitats.

#### 4.2.2 Land use and spatial correlates of feeding

Compared to the other *Culex* species *Cx. quinquefasciatus* appears to have a strong preference for chickens (Hopken et al., 2021; Kading et al., 2013) and this preference declines as urbanisation increases (because chickens are less available in city centres). Conversely, we found a large inverse relationship of *Cx. pipiens* f. *pipiens* feeding on *Turdus* (thrush species) as we move into rural and forested areas (PC1 – **Figure 3**). Thrushes such as the American robin are common in suburban parks and gardens where they have high reproductive success and survival (Evans et al., 2015; Morneau et al., 1995) and closer inspection of studies revealed that *Cx. pipiens* f. *pipiens* collected from urban and suburban properties had a high proportion of these birds (Kothera et al., 2020).

Mean annual temperature, which spatially covaries with distance to the equator, was the only climate variable that had an effect on feeding patterns. It was inversely correlated with *Culex* feeding on species from the thrush genera *Turdus*. Closer inspection of the studies revealed that this was because feeding occurred in more seasonal geographic regions (with a lower mean annual temperature) and sampling was predominantly conducted during the summer when the birds were present (Hamer et al., 2009; Komar et al., 2018; Kothera et al., 2020). Unfortunately, the studies we analysed often pooled data across dates, and we were not able to more precisely model mosquito feeding behaviour as a function of season.

### 4.3 Areas for future method/data developments

This study was an opportunity to explore what data is available and how it can be incorporated into a model, and although there are limitations, it identifies areas that need to be addressed going forward. A key feature of this study is that it replaced the environmental metadata of locally conducted mosquito diet studies with a global metadata. The climate and land use data used were collected at a 0.01-degree resolution (∼1.1km) and averaged within a 10 km radius of each sample site. While this is highly resolved at the global scale, it is relatively coarse if we consider the scale that mosquitoes generally live at (Martínez-de la Puente et al., 2020; Moore & Brown, 2022) even though many mosquitoes disperse 10-100 kilometres (Huestis et al., 2019). The land use and average climate data was centred around 2010, which is near the midpoint of the time when the studies were collected (since 2000). For domestic livestock density we were able to use the data and projections of Gilbert et al. (2018) and the GPW4 for human density (Center for International Earth Science Information Network, 2018) but this was not available for wild animal and pet densities as predictor variables.

It is unfortunate that seasonality or rapid scale changes in land use could not be included in our models. These impact host availability because of migration, natural succession or habitat suitability. For mosquitoes can seasonally switch from birds to mammals in cooler months (first observed in (Tempelis et al., 1965). Season and local landscape features also impact mosquito physiology and behaviour, with host seeking behaviour being impacted by plant availability (nutritious sugar source), rainfall, temperature and daylight hours (Carvajal-Lago et al., 2021; Farajollahi et al., 2011).

With more studies and more systematically reported metadata to become available, it would be also desirable to account from small-scale study details, including for example indoor/outdoor sampling (see supplementary figure S3), the proximity of livestock farms and zoos and local-scale habitat features.

#### 4.3.2 Omitted studies

We omitted many studies because of our strict inclusion criteria. Notably, our literature search returned 1,495 publications since 2000, which were manually checked and only 88 met our criteria. Most exclusions were because of our criteria that studies use ‘universal’ molecular methods to detect hosts. The biggest impact of this was on *Anopheles funestus* and *Anopheles gambiae* for which we recovered only 14 studies that used universal methods but a considerable number of studies used ELISA (n=44), highly targeted PCR methods (n=9) and a precipitin study. The loss of so many studies was offset by the benefit that methods which don’t make *a priori* assumptions about host range afford us a deeper and less biased insight into vector ecology, and gave greater prominence to the rare times that the *Anopheles* fed on unexpected hosts (Diallo et al., 2013; Ogola et al., 2017; Omondi et al., 2015). Some studies were also excluded because they did not systematically report the sampling locations. In many studies molecular feeding data is frequently pooled across locations or timepoints, which means local environmental or seasonal features are lost. We included these studies, to ensure that we had sufficient data to run our model, while exploring finer spatial and temporal resolution requires more refined data and awaits further research.

### 4.4 Wider implications of research and conclusion

From an epidemiological perspective, vector-borne pathogen transmission between different host species is the outcome of several features, including: host and vector range, host and vector competence, and host selection plasticity (Campos et al., 2023; Mordecai et al., 2019; Takken & Verhulst, 2013). Mosquitos that feed on diverse hosts present the greatest risk for transmitting competent pathogens between host species. However, this can limit the spread of highly specialised pathogens. For example, *Culex* mosquitoes may successfully transmit West Nile virus between competent bird species, but blood feeding on mammals as additional hosts may dilute the transmission of this virus as viral loads in mammals are typically too low for transmission (Hamer et al., 2009). If vectors and pathogens specialise in the same host species, then this can also increase transmission (Fikrig & Harrington, 2021). Intermediate feeding patterns should favour transmission of multihost pathogens. Thus, a better understanding of the drivers that cause plasticity in feeding patterns of mosquitos are crucial for better predicting pathogen transmission risks in natural environments and improve the targeting of control efforts.

## 5 Conclusion

We found that the focus on molecular studies that use ‘universal’ methods considerably increased the discovery rate of novel hosts per sample measured. This is mostly because although mosquitoes generally feed on the hosts that they are expected to – there are also many anomalies (e.g. mammalophilic mosquitoes feeding on birds or reptiles). Our meta-analysis set out to understand what we can learn by putting local blood meal studies in a global context. We confirm that the host selection is not solely determined by innate preference and host availability but also by other environmental features. While our strict criteria for inclusion meant that only 88 out of 1,495 studies that we read were included in the analysis, these results suggest that universal molecular methods are a means toward understanding how environment shapes mosquito blood feeding and the corresponding risks of disease transmission and spillover.

## DATA AND CODE ACCESSIBILITY

Data and code are accessible from the following GitHub repository: https://github.com/konswells1/Feeding-patterns

## ACKNOWLEDGMENTS

K.W. and T.U.W. were supported by the Royal Society Grant RGS\R2\222152, N.C. was supported by the ARC-DECRA Fellowship DE210101439. M.L. acknowledges support from the James McCune Smith scholarship programme at Glasgow University. Heather Ferguson and Kimberly Fornace (Glasgow University) provided valuable feedback on a draft.

## AUTHOR’S CONTRIBUTIONS

K.W. conceived first ideas and helped M.L. explore preliminary data as part of his BSc dissertation at Swansea University; R.O’R. led the subsequent data compilation with help from K.W., T.U.W, and M.L; K.W. and N.C. analysed the data; K.W. wrote the first draft. All authors contributed to discussions, interpretations, manuscript writing, and gave final approval for publication.

