## Supplementary materials for "The influence of environment on mosquito feeding patterns: a meta-analysis of ‘universal’ DNA diet studies in a global context"

**Supplement content**

| Item | Description |
|---|---|
| Figure S1 | Global map of study sites |
| Figure S2 | Correlation plot among 15 environmental variables extracted for all study sites |
| Figure S3 | Correlation plot among the nine environmental predictor variables used in the analysis |
| Figure S4 | Relative effects of different environmental predictors on the shifts in mosquito blood meal origin from different taxonomic host groups |
| Table S1 | Overview of the number of studies retrieved from Web of Science using complementary search strategies |
| Table S2 | Description of environmental variables used to characterize study sites |

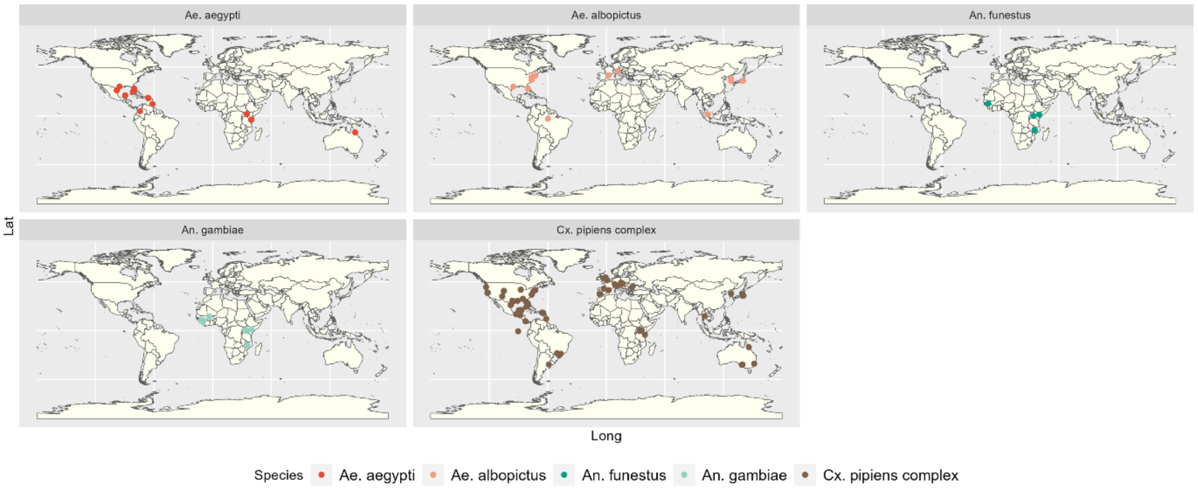

**Figure S1.** Study sites at which data on blood meal origin from the six focal mosquito species were available. These were *Aedes albopictus*, *Aedes aegypti*, *Anopheles gambiae*, *Anopheles funestus*, and *Culex pipiens* species complex. The maps show that we have reasonable sampling coverage across the geographic range of these taxa.

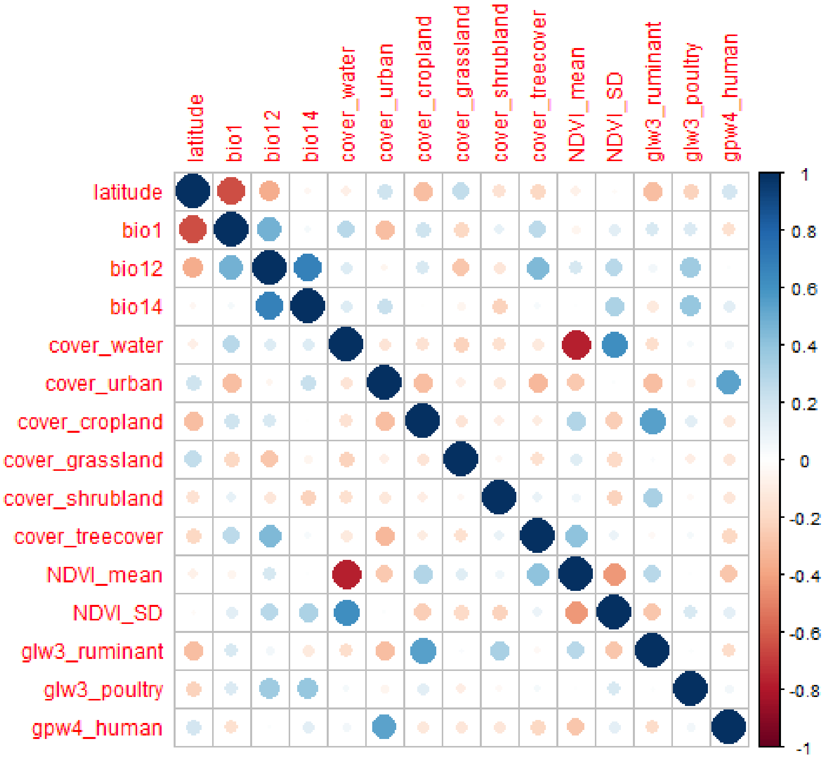

**Figure S2.** Correlation plot among 15 environmental variables extracted for all study sites prior to selection of variable with minimal pair-wise correlation for modelling.

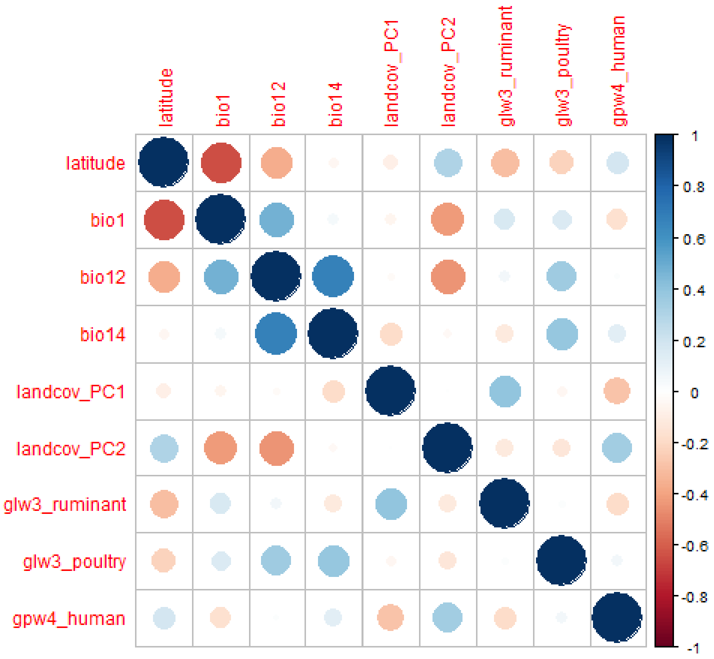

**Figure S3.** Correlation plot among the nine environmental predictor variables used in the analysis.

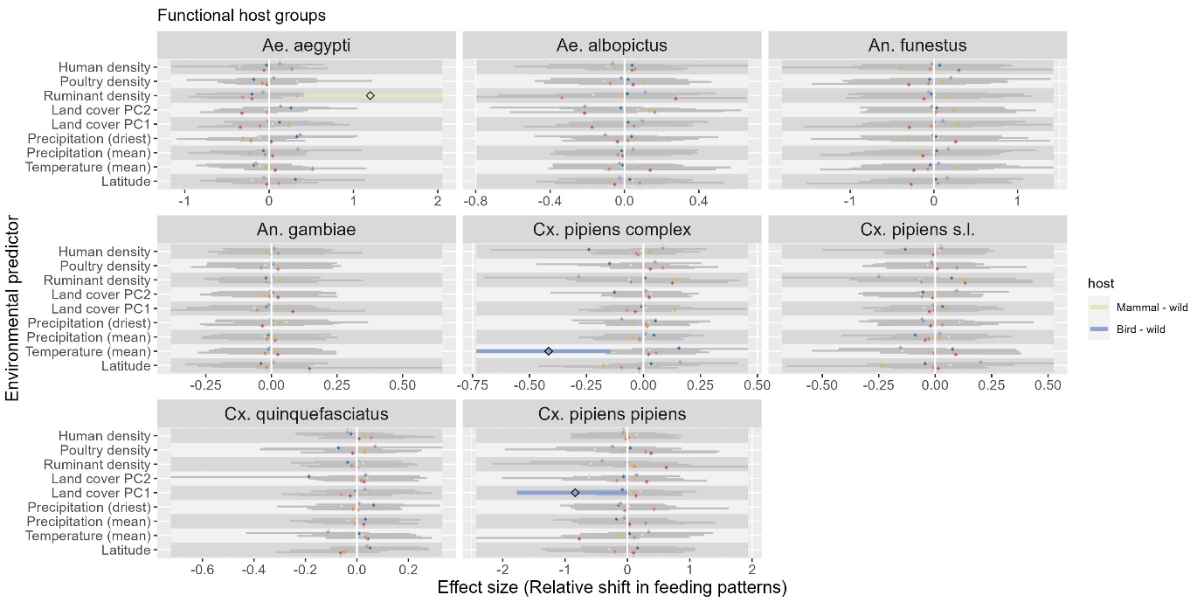

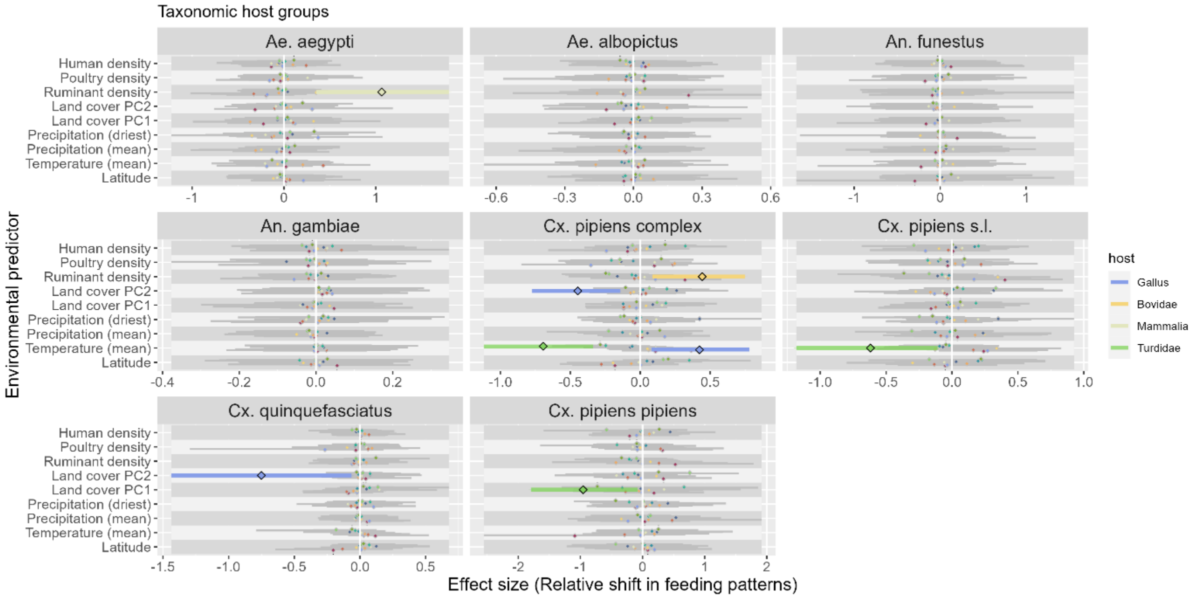

**Figure S4.** Relative effects of different environmental predictors on the shifts in mosquito blood meal origin from different functional (panel A) and taxonomic (panel B) host groups for a subset of studies excluding those that captures mosquitoes indoors only (see main text for equivalent plots for the full data set analysis). Points and error bars represent the scaled coefficient estimates (mean and 90% CI credible intervals) from models of variation in relative host utilisation frequencies for humans (red), mammalian pets (blue), mammalian wildlife (green) and wild birds (dark blue), whereby only estimates not overlapping one are displayed in thick-lined colour and those overlapping one in thin-lined grey (including all host groups for which no relationship were evidenced). Abbreviated genera names correspond to *Aedes, Anopheles and Culex*. *Cx. pipiens* s.l. and *Cx. quinquefasciatus* samples were also combined into *Cx pipiens complex* and we also conducted analysis for *Cx. pipiens* f. *pipiens*. Note that with only a single outdoor study available for *Cx. pipiens* f. *molestus*, analysis on the data subset was not conducted for this biotype.

**Table S1** Overview of the number of studies retrieved from Web of Science using complementary search strategies. The number of studies for each row making the final analysis are given in the "Final Set" column whereas the columns entitled "Web Science" and "AND [methods]"/ "AND 'mosquito*'" indicates that numbers of studies return from the respective search strings (with and without filtering for methods/mosquitoes, respectively). To gauge search term redundancy, the "unique" column records if any publications in the final set do not occur in any other search row. The different search strategies are represented in the table as follows: **A** Blood-meal analyses publications were identified using the search terms "'bloodmeal*' OR 'blood meal' OR 'blood-meal'" (denoted as 'blood*' in table) along with a mosquito identifier. If there were too many irrelevant results, then a disjunctive list of blood-meal analysis methods were added to the search. **B** Popular vertebrate, avian and mammalian PCR primer sets were also used as search terms. If there were too many irrelevant results, then the search was narrowed by searching for the phrase 'mosquito*' in the retrieved literature. **C** To assess how comprehensive the literature search was, we searched literature that we have personally stored on our personal bibliographies over the past few years.

| **1A Search by vector name** | **Web of Science** | **AND [methods]** | **Final Set** | **Unique** |
|---|---|---|---|---|
| 'blood*' AND 'mosquito*' | 2980 | 528 | 39 | 1 |
| 'blood*' AND *'Culex pipiens'* | 314 | 58 | 17 | 0 |
| 'blood*' AND '*Culex quinquefasciatus*' | 260 | 53 | 15 | 0 |
| 'blood*' AND '*Culex pipiens quinquefasciatus'* | 68 | - | 7 | 0 |
| 'blood*' AND '*Culex pipiens fatigans'* | 3 | - | 0 | 0 |
| 'blood*' AND *'Culex p fatigans'* | 4 | - | 0 | 0 |
| 'blood*' AND '*Culex p pallens'* | 8 | - | 0 | 0 |
| 'blood*' AND '*Culex pipiens pallens'* | 22 | - | 3 | 0 |
| 'blood*' AND '*Culex p molestus'* | 31 | - | 4 | 0 |
| blood_* 'AND '*Culex pipiens molestus'* | 17 | - | 0 | 0 |
| 'blood*' AND *'Aedes aegypti'* | 1383 | 157 | 7 | 0 |
| 'blood*' AND *'Aedes albopictus'* | 344 | 66 | 9 | 0 |
| 'blood*' AND *'Anopheles funestus'* | 123 | 61 | 3 | 0 |
| 'blood*' AND *'Anopheles gambiae'* | 815 | 191 | 4 | 0 |
| **1B Search by methods publication** | **Web of Science** | **AND 'mosquito*'** | **Final Set** | **Unique** |
| Abbasi *et al* 2009 | 55 | 1 | 1 | 0 |
| Alcaide et al 2009 | 112 | - | 17 | 0 |
| Berdjane-Brouk *et al* 2012 | 58 | - | 0 | 0 |
| Boakye *et al* 1999 | 132 | 46 | 11 | 0 |
| Cadenas 2007 | 78 | 3 | 0 | 0 |
| Cicero and Johnson 2001 | 92 | 20 | 8 | 0 |

| | | | | |
|---|---|---|---|---|
| Egizi *et al* 2013 | 22 | - | 6 | 0 |
| Field *et al* 2019 | 3 | - | 0 | 0 |
| Fornadel and Norris 2008 | 34 | - | 4 | 0 |
| Humair *et al* 2007 | 116 | 17 | 0 | 0 |
| Ivanova *et al* 2007 | 396 | 12 | 6 | 1 |
| Kent 2009 | 173 | - | 25 | 1 |
| Kitano *et al* 2007 | 162 | 25 | 4 | 2 |
| Kocher *et al* 1989 | 4144 | 63 | 10 | 0 |
| Meece et al. 2005 | 29 | - | 4 | 0 |
| Melton *et al* 2007 | 51 | 1 | 28 | 0 |
| Molaei et al. 2006 | 269 | - | 26 | 3 |
| Ngo and Kramer 2003 | 137 | - | 18 | 0 |
| Omondi et al 2015 | 33 | 17 | 4 | 0 |
| Purdue *et al* 2000 | 60 | - | 1 | 0 |
| Reeves et al 2016 | 26 | - | 0 | 0 |
| Reeves *et al* 2018 | - | - | 7 | 1 |
| Sawabe et al 2010 | 55 | - | 10 | 1 |
| Townzen et al 2008 | 87 | - | 6 | 0 |
| Roca et al 2004 | 107 | 5 | 0 | 0 |
| **1C** | **Publications on researchers' hard drives** | **Final Set** | **Unique** | |
| External gauge on meta-analysis completeness | 470 | 84 | 12 | |

ϕ 'Precipitin' OR 'ELISA' OR 'PCR' or 'amplicon' OR 'metabarcod*' OR 'MALDI'

**Table S2.** Description of environmental variables used to characterize study sites where blood meals of mosquitos have been recorded. Only those marked with an asterix and grey-shaded rows have been considered for analyses.

| **Variable** | **Description/ Source** |
|---|---|
| Annual mean temperature* | Annual mean temperature (bio1) records from WorldClim version 2.0 |
| Annual mean temperature* | Annual mean temperature (bio12) records from WorldClim version 2.0 |
| Rainfall of driest month* | Rainfall of driest month (bio14) records from WorldClim version 2.0 |
| Precipitation seasonality | Precipitation seasonality (bio15) records based on the coefficient of variation, from WorldClim version 2.0 |
| Elevation | Elevation, extracted from SRTM 90m Digital Elevation Database v4.1 |
| Latitude* | Latitude as given by geographical coordinate |
| Cropland cover | Proportional landcover within 10km radii around study site locations; Copernicus landcover data version 2.07 from 2010 |
| Water bodies cover | Proportional landcover within 10km radii around study site locations; Copernicus landcover data version 2.07 from 2010 |
| Urban cover | Proportional landcover within 10km radii around study site locations; Copernicus landcover data version 2.07 from 2010 |
| Grassland cover | Proportional landcover within 10km radii around study site locations; Copernicus landcover data version 2.07 from 2010 |
| Tree cover | Proportional landcover within 10km radii around study site locations; Copernicus landcover data version 2.07 from 2010 |
| Shrubland cover | Proportional landcover within 10km radii around study site locations; Copernicus landcover data version 2.07 from 2010 |
| NDVI (mean) | Mean of the normalized difference vegetation index (NDVI) for the year 2010 in buffers of 10 km radius around all sampling locations; Terra Moderate Resolution Imaging Spectroradiometer (MODIS, MOD13Q1 version 6) |
| NDVI (SD) | SD of the normalized difference vegetation index (NDVI) for the year 2010 in buffers of 10 km radius around all sampling locations; Terra Moderate Resolution Imaging Spectroradiometer (MODIS, MOD13Q1 version 6) |
| Human density* | Human population density based on Gridded Population of the World version 4.0 dataset |
| Buffalo density | Density of domestic buffalo in 10km radius around study sites based on Gridded Livestock of the World (GLW3) database |
| Cattle density | Density of domestic cattle in 10km radius around study sites based on Gridded Livestock of the World (GLW3) database |
| Goat density | Density of domestic goat in 10km radius around study sites based on Gridded Livestock of the World (GLW3) database |
| Horse density | Density of domestic horses in 10 km radius around study sites based on Gridded Livestock of the World (GLW3) database |
| Pig density | Density of domestic pig in 10km radius around study sites based on Gridded Livestock of the World (GLW3) database |
| Chicken density | Density of domestic chicken in 10km radius around study sites based on Gridded Livestock of the World (GLW3) database |
| Duck density | Density of domestic duck in 10km radius around study sites based on Gridded Livestock of the World (GLW3) database |
| Sheep and goat | Density of domestic sheep and goat in 10km radius around study sites based on Gridded Livestock of the World (GLW3) database |

| | |
|---|---|
| Ruminant density* | Density of domestic ruminants (sheep, goat, cattle, and buffalo) in 10km radius around study sites based on Gridded Livestock of the World (GLW3) database |
| Poultry density * | Density of domestic ruminants (chicken and dug) in 10km radius around study sites based on Gridded Livestock of the World (GLW3) database |
| Land cover PC-1 ('green spaces') * | First component from a principal component analysis of all the land cover and NDVI variables mentioned above. Increasing values represent mainly an increase in vegetated land cover types and NDVI and decrease in water bodies and urban land cover, explaining 30.5% of the variation in land cover variables across sites |
| Land cover PC-2 ('urban spaces') * | First component from a principal component analysis of all the land cover and NDVI variables mentioned above. Increasing values represent mainly an increase in urban land cover and decrease in tree cover, explaining 18.8% of the variation in land cover variables across sites. |

**Table S3.** Selected features of the principal component analysis (PCA) conducted on land cover variables (as described in Table S1 and methods in the main text).

Importance of components:

| | PC1 | PC2 | PC3 | PC4 | PC5 | PC6 | PC7 | PC8 |
|---|---|---|---|---|---|---|---|---|
| Eigenvalue | 2.4466 | 1.5054 | 1.0932 | 1.0726 | 0.9942 | 0.46925 | 0.34215 | 0.076579 |
| Proportion Explained | 0.3058 | 0.1882 | 0.1367 | 0.1341 | 0.1243 | 0.05866 | 0.04277 | 0.009572 |
| Cumulative Proportion | 0.3058 | 0.4940 | 0.6307 | 0.7647 | 0.8890 | 0.94766 | 0.99043 | 1.000000 |

Variable scores:

| | PC1 | PC2 | PC3 | PC4 | PC5 | PC6 |
|---|---|---|---|---|---|---|
| Cropland cover | 0.8383 | -0.61288 | -0.9374 | -0.76527 | -1.03978 | 0.5309 |
| Water cover | -1.7789 | -0.62121 | -0.3673 | 0.28687 | -0.19177 | -0.2569 |
| Urban cover | -0.3055 | 1.50732 | 0.9023 | -0.68856 | 0.04282 | 0.3219 |
| Grassland cover | 0.4365 | 0.55507 | -1.1968 | 0.60904 | 1.24534 | 0.3383 |
| Tree cover | 0.6725 | -1.42783 | 0.8148 | 0.05016 | 0.62972 | -0.2151 |

| | | | | | | |
|---|---|---|---|---|---|---|
| Shrubland cover | 0.4885 | 0.09081 | 0.5505 | 1.63629 | -0.75266 | 0.5203 |
| NDVI - mean | 1.8145 | -0.30642 | 0.3260 | -0.30660 | 0.36810 | 0.1644 |
| NDVI - SD | -1.3569 | -0.80976 | 0.3523 | -0.29402 | 0.53338 | 1.0017 |
